# Adherence to minimal experimental requirements for defining extracellular vesicles and their functions: a systematic review

**DOI:** 10.1101/2021.04.23.441160

**Authors:** Rodolphe Poupardin, Martin Wolf, Dirk Strunk

**Affiliations:** Cell Therapy Institute, Spinal Cord Injury and Tissue Regeneration Center Salzburg (SCI-TReCS), Paracelsus Medical University (PMU), Salzburg, Austria

**Keywords:** MISEV, extracellular vesicles (EVs), exosomes, text mining, data science, regenerative medicine

## Abstract

Rigorous measures are required to cope with the advance of extracellular vesicle (EV) research, from 183 to 2,309 studies/year, between 2012-2020. The ‘MISEV’ guidelines requested standardizing methods, thereby assuring and improving of EV research quality. We investigated how EV research improved over time.

We conducted a keyword search in 5,093 accessible publications over the period 2012-2020 and analyzed the methodology used for EV isolation and characterization. We found a significant improvement over the years particularly regarding EV characterization where recent papers used a higher number of methods and EV markers to check for quantity and purity. Interestingly, we also found that EV papers using more methods and EV markers were cited more frequently cited. Papers citing MISEV criteria were more prone to use a higher number of characterization methods.

We therefore established a concise checklist summarizing MISEV criteria to support EV researchers towards reaching the highest standards in the field.

Graphical abstract

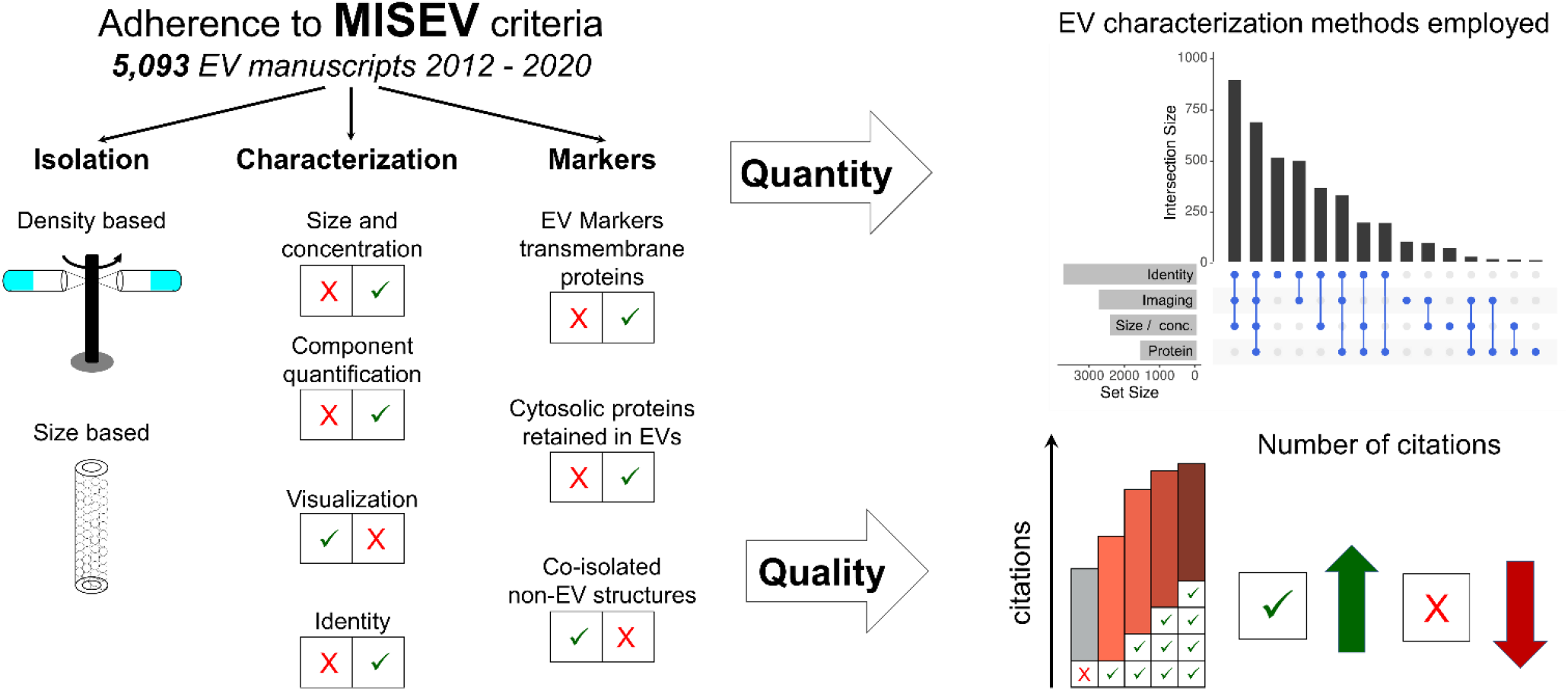

## 1. Introduction

Extracellular vesicles (EVs) are nanometer-sized membrane-enclosed particles secreted via several mechanisms by virtually all cells. They are thought to be loaded with lipids, proteins, and various nucleic acid species of the source cell [3,4]. In addition they can also carry extra-vesicular proteins accumulated from their surroundings [5]. EVs are explored increasingly as potential therapeutic and diagnostic tools due to their capacity to transfer bioactive components to recipient cells and tissues, and their contribution to intercellular communication [6,7]. EVs can support tissue regeneration, participate in immune modulation, and contribute to the mode of action of cellular therapies. Bioengineered EVs can also act as delivery vehicles for therapeutic agents [8]. The EV field has achieved widespread interest as evidenced by the increasing number of EV publications over the last years. Due to their physical nature, EVs are close to or below the detection limit of many conventional analysis methods making their investigation more challenging. Nano-sized lipoproteins or protein aggregates have overlapping characteristics with EVs concerning size or density, making proper selection of appropriate EV purification and characterization methods a key to draw concise conclusions [9,10]. The advancement of more rigorous EV research is a challenging process as EV experts are still concerned about conclusions not sufficiently supported by the information reported [11,12]. Therapeutic efficacy depends on a variety of additional parameters, which should be tested in well‐standardized quantitative assays, before being addressed in clinical trials [13]. Challenges of appropriately isolating and characterizing EV fractions and their contaminants relate to the different identity markers and their respective fractions/locations with an overlap between EVs and other contaminants regarding identity, density and size (**Figure 1;** modified from [14]). Accordingly, combinations of isolation methods and characterization categories need to be implemented for improving EV preparations and conclusions drawn from the respective experiments.

**Fig. 1:**
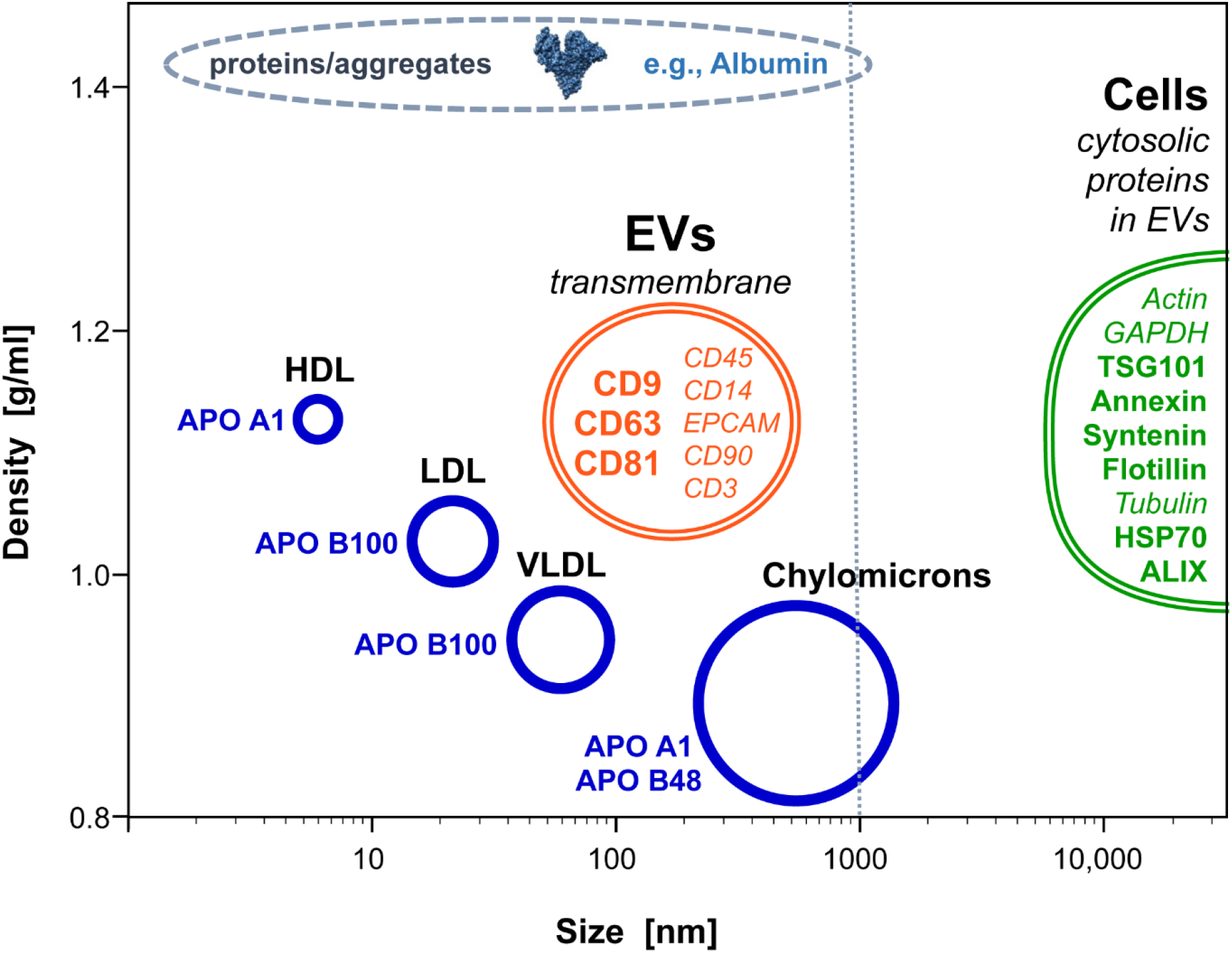
Size and density distribution of extracellular vesicles (EVs) and common contaminating particles and their molecular markers. Diagram showing the overlap between different lipoprotein fractions (blue), extracellular vesicles (orange), protein aggregates (grey) and cells (green) in size (x axis; logarithmic) and density (y-axis). In addition, representative molecular markers found to be most used in our analysis and which were recommended in MISEV2018 as universal markers for identification of extracellular vesicles (orange, bold). Transmembrane markers of extracellular vesicles derived from specific cell lineages are shown in *italic* type. Cytosolic proteins recovered inside vesicles and related to vesicle biogenesis are shown in bold green while cytosolic proteins that are frequently retained in vesicles are shown in *italics*, green. Markers of specific lipoproteins potentially co-isolated with EVs are shown in blue. The structure of albumin (PDB: 1E78) representing the most common protein contamination was rendered using UCSF Chimera [16]. Figure adapted from [14].

In 2014, the International Society for Extracellular Vesicles (ISEV) proposed comprehensive Minimal Information for Studies of Extracellular Vesicles (MISEV) guidelines suggesting protocols and steps to follow for documenting specific EV-associated functional activities [1]. In 2018, these guidelines were updated [2] and both manuscripts received robust interest in the community as shown by their high number of citations (cited more than 1,400 and 1,700 times, respectively). In addition, the data depository platform EV-track (https://www.evtrack.org) was created to promote better standardization and to provide higher research quality and transparency in the field regarding the isolation and characterization of EVs [15]. Researchers have been invited to make their complete method details available to EV-track on a voluntary basis. This platform provides valuable information on the MISEV2018 criteria adherence of the manuscripts. Because of its nonobligatory nature potentially leading to increased bias, EV-track may not completely reflect the global state and quality of the EV field.

In addition, surveys were conducted to assess the methods used by researchers and how trends are evolving [4,17]. In 2015, 196 responses were received, and 320 full responses plus 300 partial responses were received in 2020. Despite a potential bias due to the “voluntary basis” of these surveys, authors demonstrated the lack of controls before and after EV separation, highlighting the importance of defining clear guidelines. Other databases such as *Exocarta* (https://www.exocarta.org) or *Vesiclepedia* (https://www.microvesicles.org) fulfill a different purpose by providing a manually curated compendium of molecular data on lipids, proteins and RNAs identified in different classes of EVs (for *Vesiclepedia*) or in exosomes (for *Exocarta*). Although all methods used to isolate and characterize the EVs are provided together with the data, their main focus has been more on the content of EVs rather than on the fulfillment of the MISEV guidelines.

In this context, we conducted an unbiased review using a text mining approach to assess adherence to MISEV criteria [1,2] and analyzed 5,093 open access EV papers published over the period of 2012 - 2020. We first determined EV isolation methods most frequently used over time and next investigated how EV identity, purity and quantity were determined. We also examined, whether publications with higher number of characterization methods and identification markers would result in a higher recognition by the scientific community, as evidenced by the number of citations.

## 2. Development of a text mining approach to assess adherence to MISEV guidelines

### 2.1 Dataset acquisition

Publications were selected in PubMed using the *RISMed* R package [18]. We searched for manuscripts containing the keywords “extracellular vesicles” or “vesicles” together with “exosomes” in the title or abstract or papers with the major Mesh terms “extracellular vesicles” or “exosomes” (without looking at the child terms (non-explosive search)): (“extracellular vesicles”[TIAB]) OR (“exosomes”[TIAB] + “vesicles”[TIAB]) OR (“extracellular vesicles”[majr:noexp)] OR (“exosomes”[majr:noexp]). A first filter step was applied on the metadata to remove conference / symposium / case reports / reviews and editorial abstracts. PDF links of open access publications were then obtained using the R package *roadoi* [19] which is a client for the *Unpaywall-API* (https://unpaywall.org/). An R script automatically downloaded 5,495 manuscripts (using the *download.file()* command with the “*wget*” method) which were sorted according to journal’s name and year of publication. We focused on the period of 2012 - 2020 as the reference journal in the field, the ‘Journal of Extracellular Vesicles’, was first published in 2012.

### 2.2 Text mining

Our approach to detect the different methods used was based on a keyword search using regular expressions (*regex*). In order to reduce the rate of false positives, we limited the search in all downloaded PDFs to the material and methods and results parts. All keyword searches were conducted using the R package *pdfsearch* [20]. The first part comprised extracting the line numbers in each PDF for the headers introduction / material and methods / results / discussion and references to define the search area. If the result part was merged with discussion, then only material and methods part was used to avoid the detection of false positive keywords as discussion may cite methods that were not used within the manuscript (see **suppl. table 1** for complete journal list and **suppl. file 2** (R code) for exact *Regex* searches). If material and methods and results were both missing, suggesting review manuscripts, then the PDF was removed from the analysis.

We sorted the keywords into sub-categories for the EV isolation and characterization parts. A more detailed analysis was conducted to detect which markers were used based on the MISEV2018 criteria [2]. To check for significant trends over time, linear regression analysis was conducted using R. Finally, we investigated whether the quality of identification and characterization of EVs in the manuscripts correlated with the number of citations. To extract the number of citations from each manuscript, the R package *rcrossref* was used [21].

## 3. The EV field is expanding at a fast pace

Searches for (“extracellular vesicles”[TIAB]) OR (“exosomes”[TIAB] + “vesicles”[TIAB]) OR (“extracellular vesicles”[majr:noexp)] OR (“Exosomes”[majr:noexp]) in PubMed retrieved 13,529 records for the period 1990 – 2020, showing that the EV field was prosperous at a fast pace leaping from 183 records in 2012 to 2,309 records/year until the end of 2020 (**suppl. Fig. 1, Fig. 2A**). For comparison, we also included the number of publications reported to the EV-track database during the observation period [15], demonstrating a spike in 2014 with 452 records for this year followed by a steady decline for the next six years, indicating limited recognition of the platform in the community. After removing manuscripts labeled as letter / conference / reviews / editorial / case reports / manuscripts with both missing material and results plus discussion parts, we reduced our dataset to 9,462 manuscripts. Out of these, we were able to download 5,495 open access articles. Finally, 5,093 manuscripts containing at least material and methods or result parts were used for the text mining analysis.

**Fig. 2:**
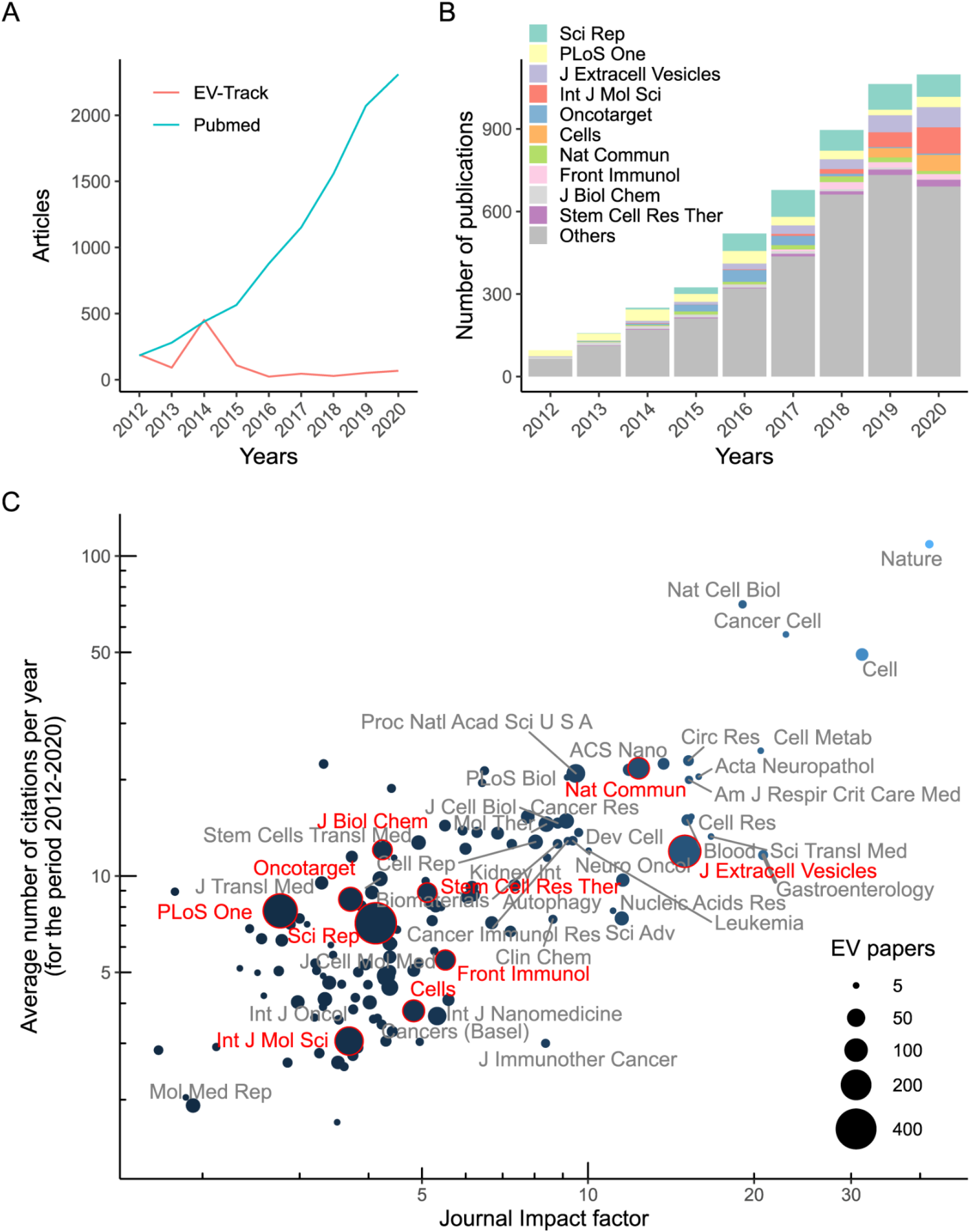
Overview of the EV publication field growth. (**A**) Steep increase of manuscripts with “extracellular vesicles” or “exosomes” + “vesicles” terms found in the title and/or abstract or with the PubMed MeSH terms “extracellular vesicles” or “exosomes” (PubMed search, 9,462 manuscripts, not considering conference / reviews / editorial / case reports). The red line shows the number of EV manuscripts submitted to the EV-track database. (**B**) Bar plot showing the year of publication for the 5,093 selected EV papers. The 10 journals with the highest number of EV related publications (top 10) are shown in different colors as indicated. (**C**) Dot plot showing the relation between journal impact factor and average number of citations for 5,093 selected EV papers between 2012 and 2020. The dot size indicates the number of EV papers found per journal. The top 10 journals identified in Fig. 2B are highlighted in red. Only journals including more than five EV papers within the observation period 2012 – 2020 are shown.

**Fig. 3:**
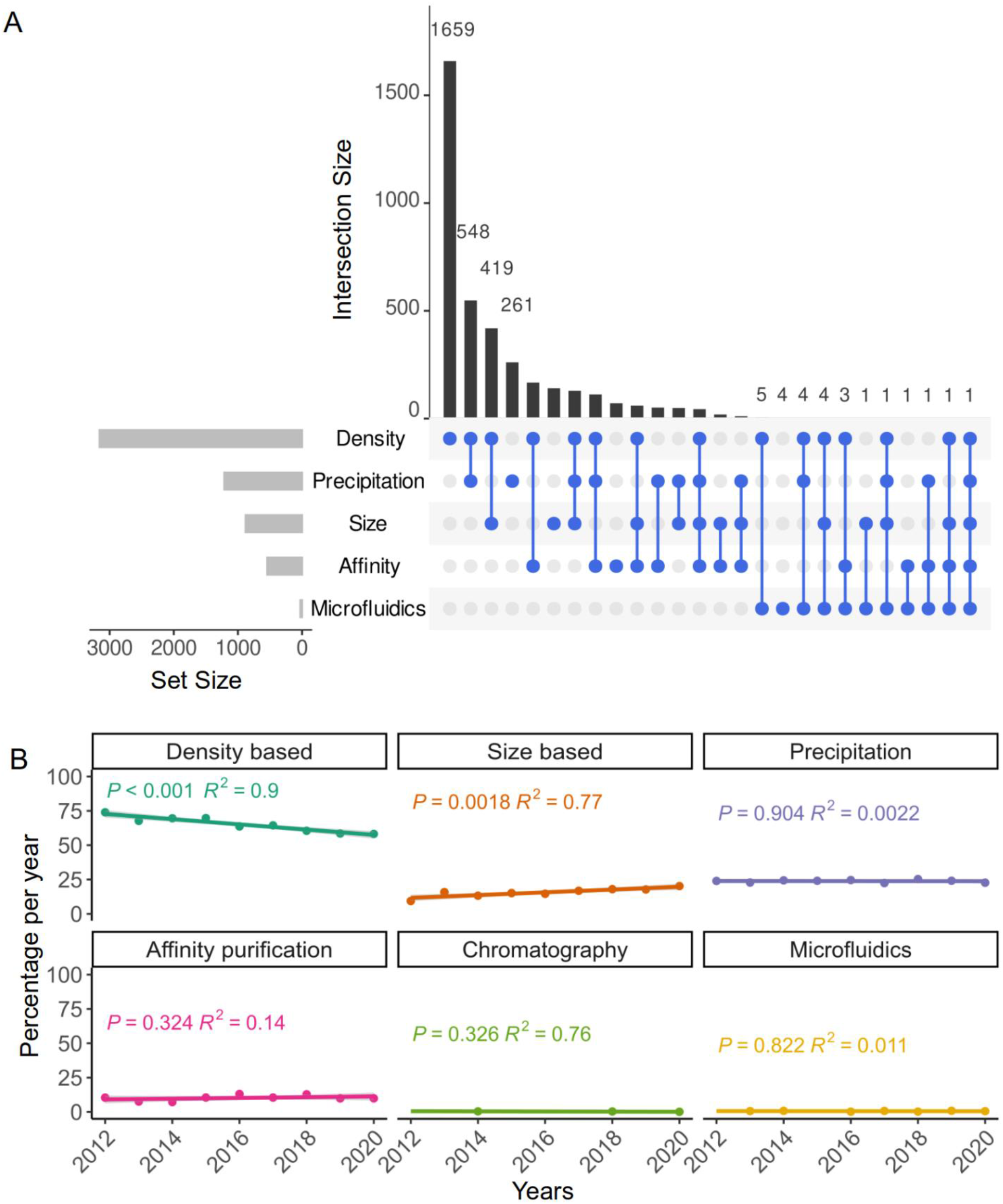
EV isolation methods. (**A**) The upset plot shows on the left the counts (set size) of the different methods ranked from the most used on the top (density-based isolation) to the least used on the bottom (microfluidics). Intersection size shows the number of publications found for each combination, ranked in descending order. The dot chart corresponding to the upper histogram shows the most used combinations of isolation methods. (**B**) Percentage of papers using the different methods during each year. Linear regression analysis identified significant changes over time (*P* < 0.05).

**Fig. 4:**
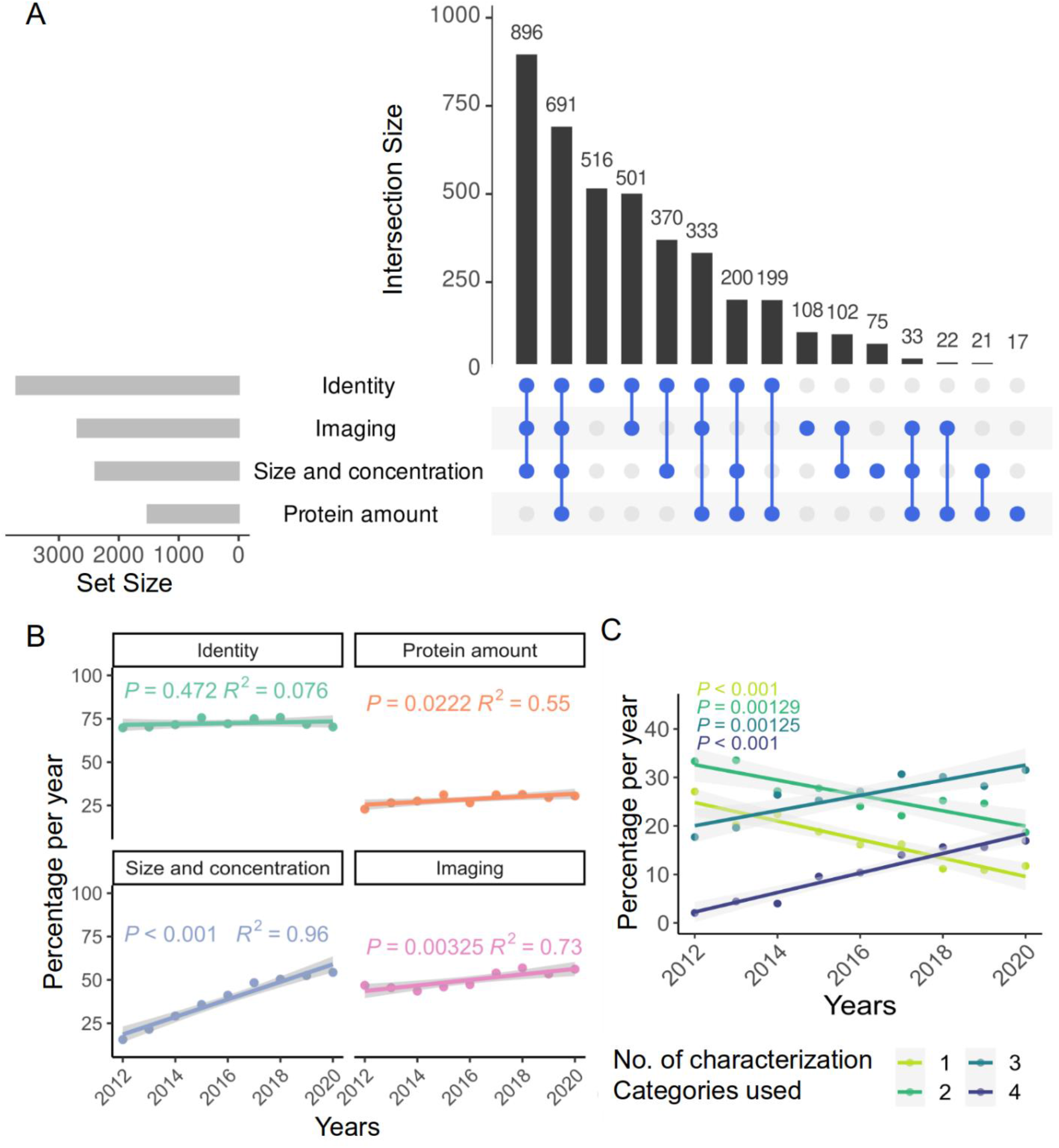
EV characterization methods. (**A**) The upset plot shows on the left the set size counts of the different methods ranked from the most used on the top (identity) to the least used on the bottom (protein amount). The upper histogram (in descending order) and the dot chart show the most used combinations for characterizing EVs. (**B**) Percentage of papers using the different methods for each year. (**C**) Percentage of EV manuscripts using a combination of characterization methods (identity, protein amount, size and concentration, imaging; color code indicated by legend). Linear regression analyses indicated significant changes over time for (B) and (C) (*P* < 0.05).

The EV manuscripts selected in our study were found in a wide range of journals. Publications from 810 journals were selected including 205 journals containing at least five EV-related manuscripts (**suppl. table 1**). Most manuscripts were found in ‘Scientific Reports’ (439), ‘PLoS One’ (284) and ‘Journal of Extracellular Vesicles’ (JEV; 247; **Fig. 2B and 2C**). The top 10 journals in our selected list are highlighted in **figure 2B**. In 2019 and 2020, a high proportion of manuscripts (244; 11.3% of total manuscripts) were published in the ‘International Journal of Molecular Sciences’ and ‘Cells’.

## 4. A broad range of methods are found for EV isolation and characterization

In order to conduct our text mining analysis, we grouped the broad range of methods for preparing and characterizing EVs in different categories based on their biophysical characteristics (**Table 1**). There is no general agreement in the field on standard method(s) for EV isolation so far. The MISEV2018 guidelines differentiate methods with various ratios of recovery and specificity [2]. Therefore, we investigated which methods or combination of methods were used and how these preferences changed over time. For EV isolation methods, our categories were based on the physical isolation principal, like ‘density-based’ for ultra-centrifugation-based enrichment strategies, or ‘size-based’ including size-exclusion chromatography and several additional filtration methods. In addition, we included ‘chemical-based’ methods like precipitation, which was applied by using commercial EV isolation kits or ‘affinity purification’ methods using antibodies or other ligands that bind specifically to EVs. Chromatography using ‘electrostatic or hydrophobic interactions’ to separate EVs from contamination substances and ‘microfluidics’ were identified among recently implemented methods.

**Table 1:**
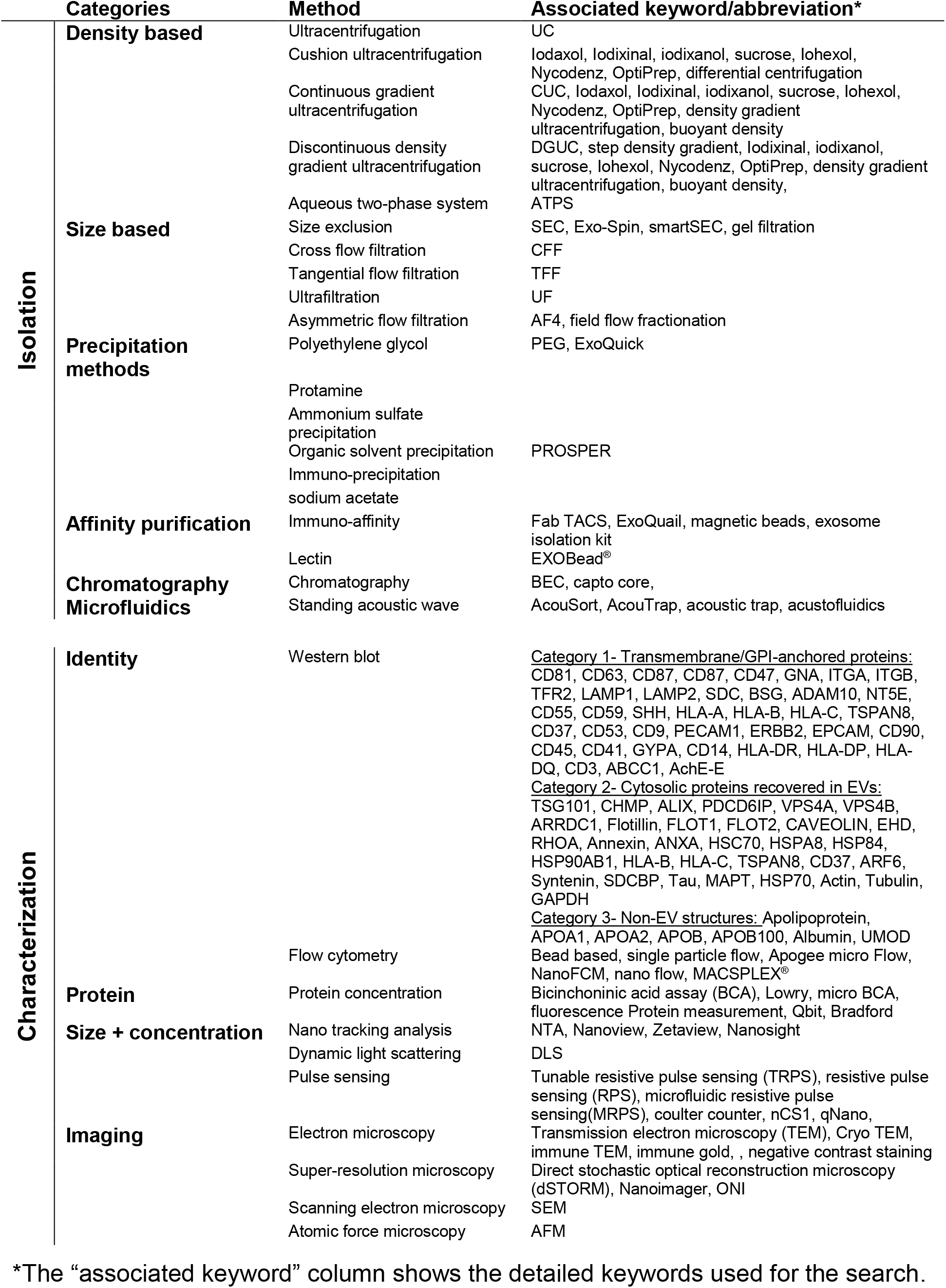
Categories and corresponding keywords used for the text mining analysis of EV manuscripts.

Concerning the categories for EV characterization methods, we adhered to the list of criteria that was recommended to evaluate EV preparations in the MISEV2018 position statement [2]. We grouped them, according to biophysical properties and the task they fulfill, i.e., checking for identity, purity or integrity. The first category, identity, was analyzed in more detail in **figure 5**. In addition to identity characterization by specific EV markers, such as tetraspanins, it is also considered important to determine the purity of EV fractions by testing for contaminants, like albumin, representing the most abundant plasma protein. The second category, protein quantification, identifies purity and determines the quantity of contaminating soluble proteins. The third category summarizes methods to evaluate EV integrity and purity by checking for size and concentration. These two points were combined as many methods give both results like nano-tracking analysis or tunable resistive pulse sensing. Finally, we summarized methods that give an actual image of the EVs illustrating round shape, morphology, double-layered membrane, size and, in some cases, also identify specific markers.

**Fig. 5:**
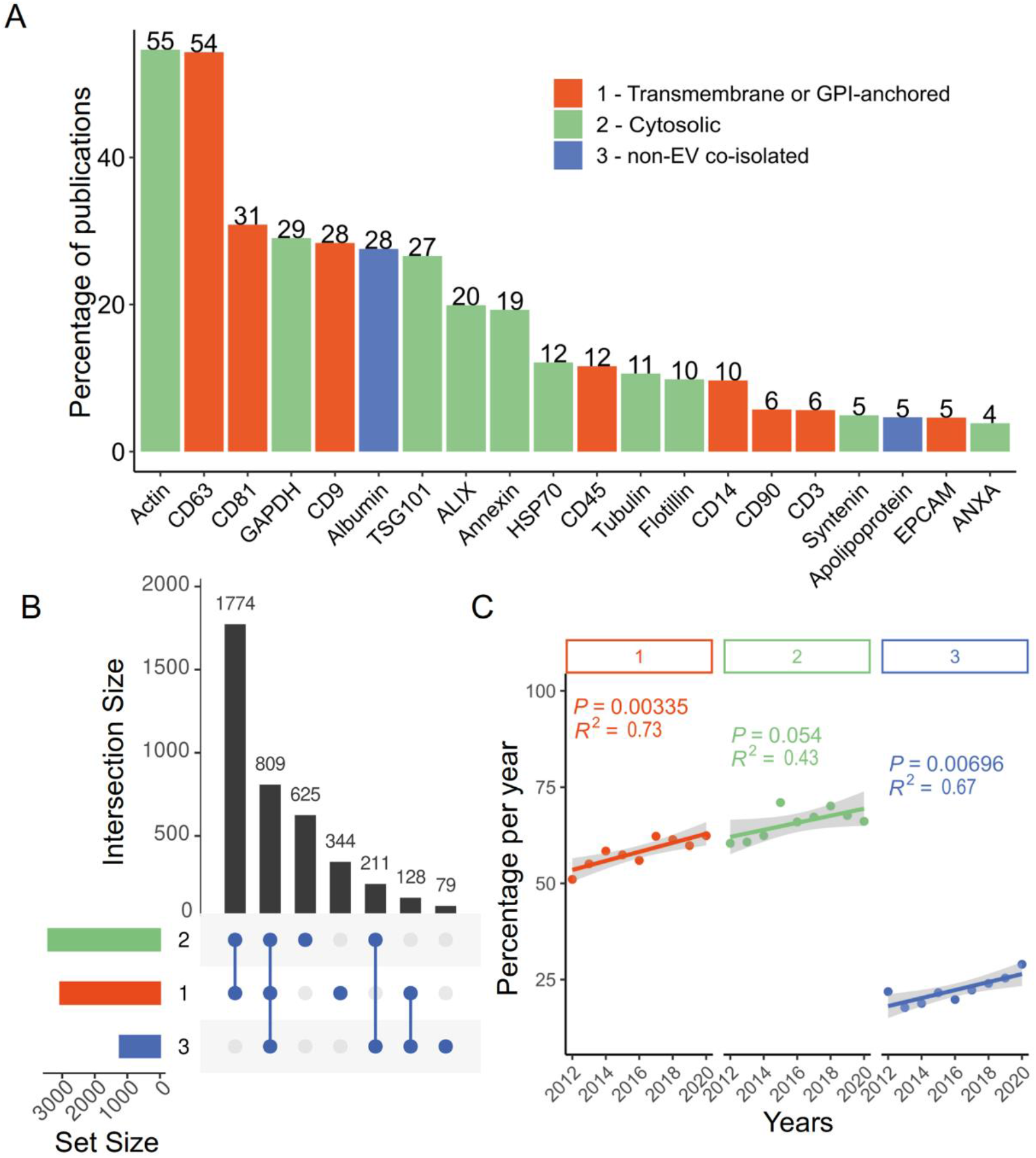
Analysis of proteins linked to EVs or contaminants. (**A**) Most used marker proteins ordered according to the frequency of use displayed with a color code indicating the category as defined by the MISEV2018 criteria (category 1: **Transmembrane or GPI-anchored** proteins associated to plasma membrane and/or endosomes; category 2: **Cytosolic** proteins recovered in EVs; category 3: Major components of **non-EV co-isolated** structures). (**B**) The upset plot shows on the left the set size counts of the different marker categories (left bars; color code as in A) ranked from the most used on the top to the least on the bottom. The upper histogram (in descending order) and the dot chart show the most used combinations of marker proteins for characterizing EVs. (**C**) Percentage of manuscripts per year using marker proteins from categories 1 - 3 to characterize their EV preparations over the years (2012 - 2020). Linear regression analyses identified significant changes over time as indicated (*P* < 0.05).

## 5. Isolation methods used for EV preparations

In our analysis of EV preparation methods, we found that most studies (1,659 / 5,093; 33.2%) used a single density-based isolation technique (**Fig. 3A**).

Other single methods identified were precipitation-based methods, used by 261 publications (5.1%) and size-based methods. The combination of density-based and precipitation methods was the second most frequently applied strategy (548 / 5,093; 10.8% of the manuscripts). This was followed by the combination of density- and sized-based methods applied by 419 studies (8.2%). Microfluidics were used in 26 papers (**Fig. 3A**). Except for size-based isolation with a significant increase over time and density-based methods showing a significant decrease, other methods’ application did not change significantly (**Fig. 3B**).

## 6. Characterization of EV fractions: the more, the better?

### 6.1 Characterization methods used

One of the key aspects of MISEV2018 [2] criteria relates to the importance of properly characterizing purified EVs regarding their identity, protein amount and physical aspects through size distribution and imaging. We found 80% of the analyzed manuscripts using at least one category. The most used characterization category was “identity”, followed by “imaging”, “size and concentration” and “protein concentration”. A higher proportion of manuscripts (17.6%) used a combination of characterization methods with “imaging”, “size and concentration” and “protein concentration”, and 13.6% used all four characterizing categories (**Fig. 4A**). A notable aspect is the significant increase of the categories, “protein amount”, “size & concentration” and “imaging” over time (**Fig. 4B**). Finally, by looking at the combination of different categories, we detected a significantly increasing percentage of manuscripts using a combination of three or four characterization methods over the years 2012 - 2020, while manuscripts using only one or two characterization methods significantly decreased over time (**Fig. 4C**).

### 6.2 Markers used for defining EV identity

Regarding EV identity, MISEV2018 [2] defined a list of protein markers divided into five groups that can be used for assessing the level of purity or contamination of the EV preparation, respectively. Researchers should use at least one marker from each of the three first categories (category 1: transmembrane or GPI-anchored proteins associated with the plasma membrane and/or endosomes; category 2: cytosolic proteins recovered in EVs; category 3: major components of non-EV co-isolated structures). The three most frequently used markers for category one were tetraspanins CD63, CD81, CD9. Actin, GAPDH and TSG101 were used most frequently for category two, and albumin as well as apolipoproteins for category three (**Fig. 5A**) We next analyzed marker combinations revealing that 34.9% of studies used a combination of category one and two, and 15.9% used the combination of three categories (**Fig.5B**). Finally, a significant increase of testing category one (transmembrane or GPI-anchored protein) and category three (non-EV co-isolated structure) marker determination in EV preparations was evident over time (**Fig. 5C**).

## 7. Impact of the MISEV guidelines on the EV community

We then asked whether manuscripts using more detailed characterization or testing more marker categories retrieved higher attention in the scientific community as evidenced by the median number of citations per year. Regression analysis showed that the median number of citations significantly increased depending on the number of characterization methods applied (*P* = 0.0002) or marker categories tested (*P* = 6.9 × 10^−6^) (**Fig. 6A**).

**Fig. 6:**
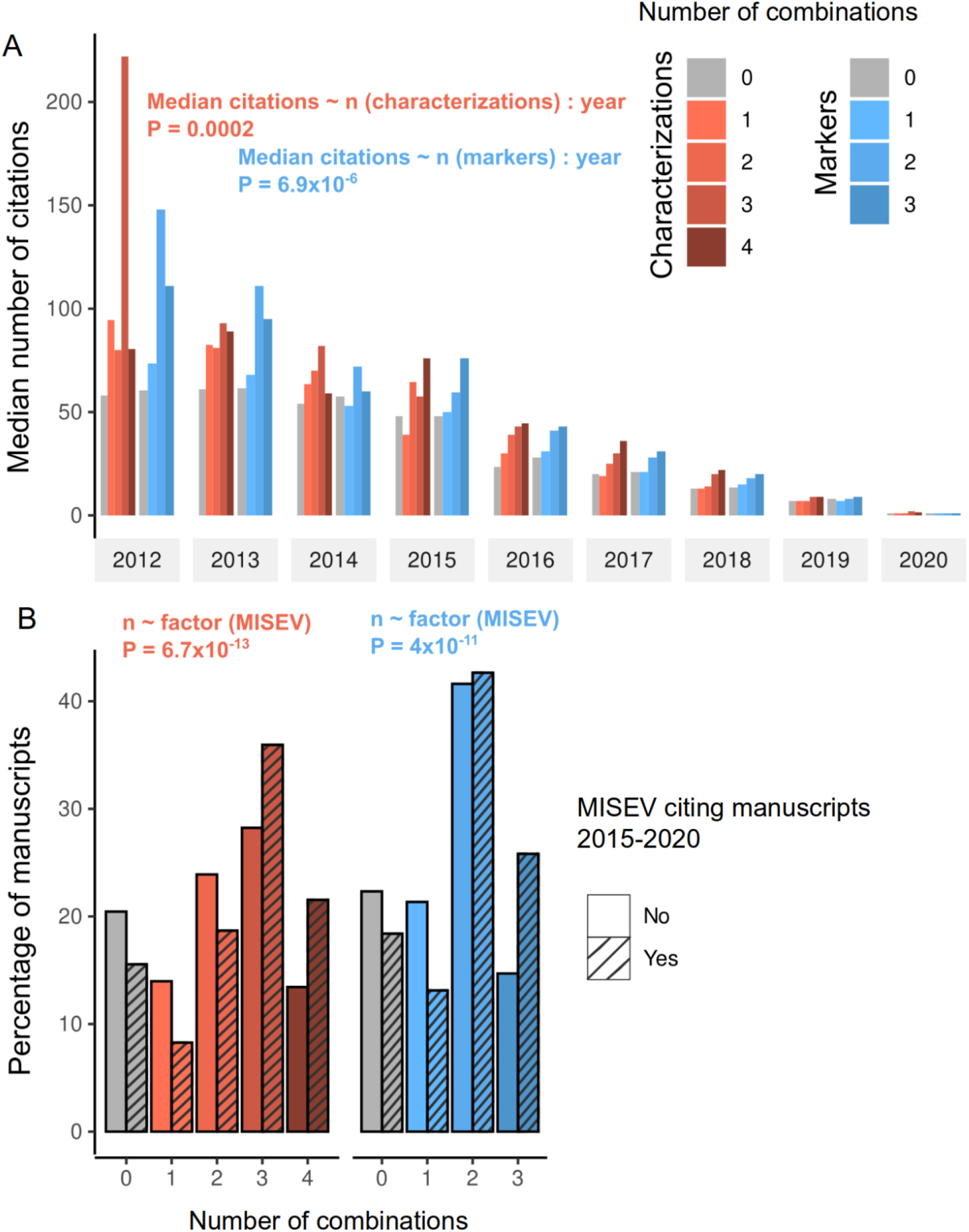
Impact of EV characterization and identification accuracy on citation frequency plus MISEV-based marker category reporting trends. (**A**) Median number of citations per year for the different combinations of characterization methods (from 0 to 4 characterization categories combined in orange-red) and marker proteins used for EV preparation characterization (0 to 3 marker categories combined (light blue-blue). Linear regression analysis revealed a significant association between the number of characterization categories and median citation numbers (*P* = 0.0002) and between the number of marker categories and median citation numbers (*P* = 6.9 × 10^−6^), respectively. (**B**) Percentage of manuscripts citing (dashed bars) or not citing MISEV (non-dashed bars) regarding the number of characterization categories used; red bars) or the different combinations of marker proteins (blue bars). Only manuscripts published between 2015 and 2020 were used for the analysis (701 manuscripts citing MISEV vs 3,878 manuscripts not citing MISEV).

By the end of 2020, MISEV2014 [1] and MISEV2018 [2] were cited more than 1,400 and 1,700 times, respectively. In order to illustrate the impact of MISEV in the EV community, we reviewed in 4,579 manuscripts, published between 2015 and 2020 and covered by our study. We found 701 manuscripts citing MISEV guidelines. We further determined the percentage of different combinations of characterization methods and markers in these manuscripts compared to the manuscripts not citing MISEV for the same period. There was a significantly higher percentage of manuscripts using 3 - 4 characterization categories (linear regression analysis, *P* = 6.7 × 10^−13^), and a lower percentage of manuscripts with no characterization in manuscripts citing MISEV, respectively, compared to manuscripts not citing MISEV. Publications citing MISEV guidelines thus appeared to be more likely to fulfill guidelines. A similar trend was found regarding the number of marker categories. There was a higher percentage of manuscripts using either two or three markers in MISEV-citing manuscripts compared to the manuscripts not citing MISEV (linear regression analysis, *P* = 4 × 10^−11^) (**Fig.6B**).

Transparent reporting of EV preparation and characterizations is an important issue. While encouraging EV researchers to submit their EV protocols to existing databases, we also developed a single-page reference table, suitable as checklist summarizing the key data and methods used for EV isolation and characterization (**Table 2**). We consider such a checklist as a tool for editors, reviewers and readers to assess the methods used and identify potentials drawbacks.

**Table 2:**
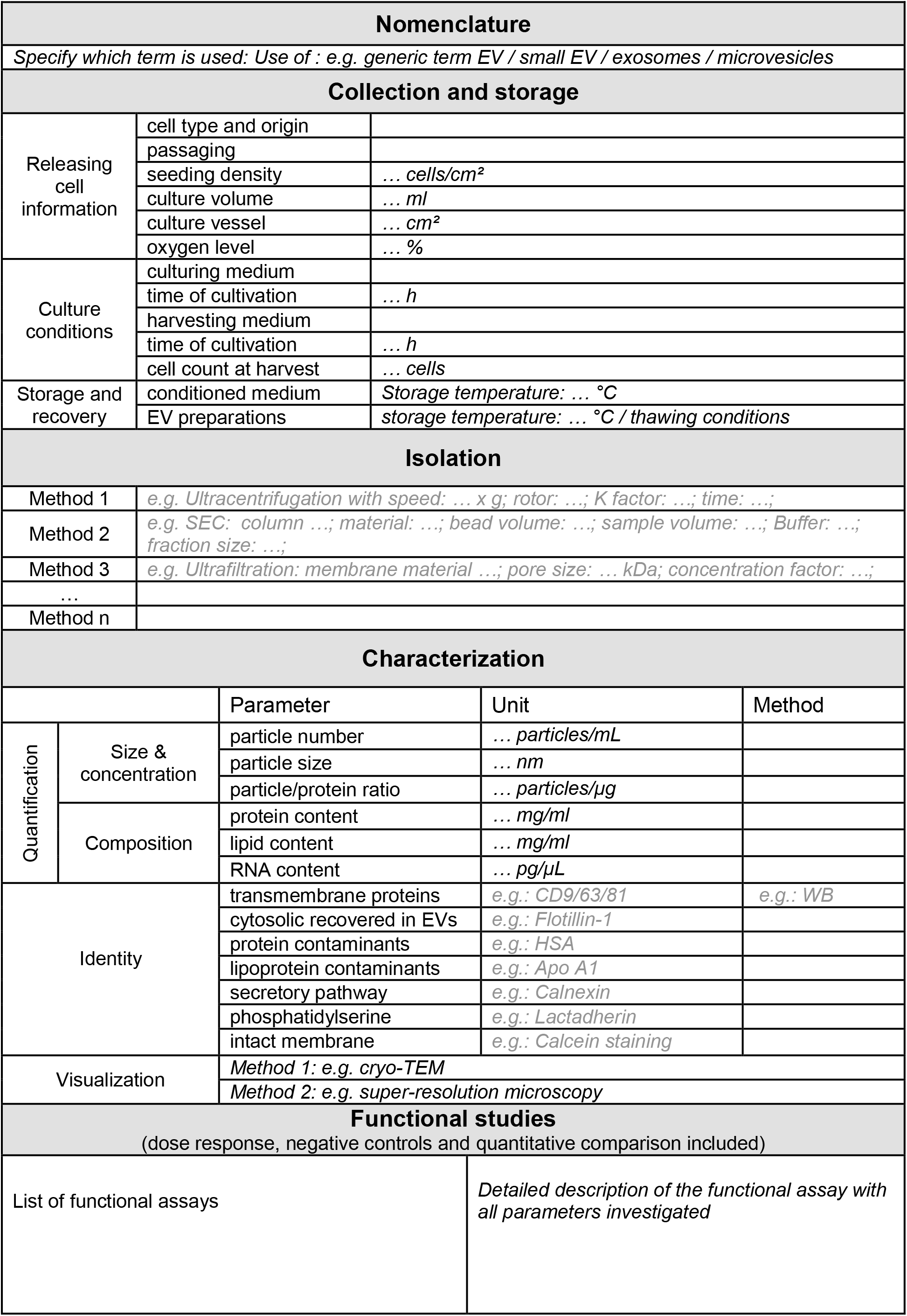
Addressing MISEV2018 criteria for research with EV material.

## 8. Conclusion

EV research is an emerging field with rapidly growing interest over the last decade. EV isolation and characterization are particularly challenging because the nano-sized cell-derived vesicles are close to or below the detection limit of many traditional methods. Additional biologically active components including lipoproteins or protein aggregates display similar size and/or density characteristics and co-purify with EVs, further confounding dedicated mechanistic studies. Recognizing these difficulties led to the establishment of the MISEV2014 and comprehensive updated MISEV2018 guidelines to set minimum standards for studies conducted with EVs [1,2]. The level of adherence to these guidelines and exploitation of additional voluntary online reporting platforms was unclear.

In this context, we conducted an unbiased review of 5,096 accessible papers published in the EV research field in a broad range of journals between 2012 and 2020. In our systematic analysis, we found that the awareness of investigators to better characterize their EV preparations using a combination of several methods was significantly rising. The majority of studies still applied only one method for EV purification with few significant changes over the past years. Studies citing the MISEV position statements used significantly more methods for EV characterization and multiple markers to determine EV identity and purity indicating the impact of the guidelines in the EV community. A precise characterization of EV preparations with multiple methods and marker categories also resulted in a significantly higher number of citations.

## Declaration of Competing Interest

The authors declare no conflict of interest in the publication of this work.

## Acknowledgements

This work was supported in part by funding from the European Union’s Horizon 2020 research and innovation program (grant agreement no. 733006 to DS), by Land Salzburg IWB/EFRE 2014 - 2020 P1812596 and WISS 2025 20102-F1900731-KZP EV-TT (to DS).

---

**Suppl. Fig. 1:**
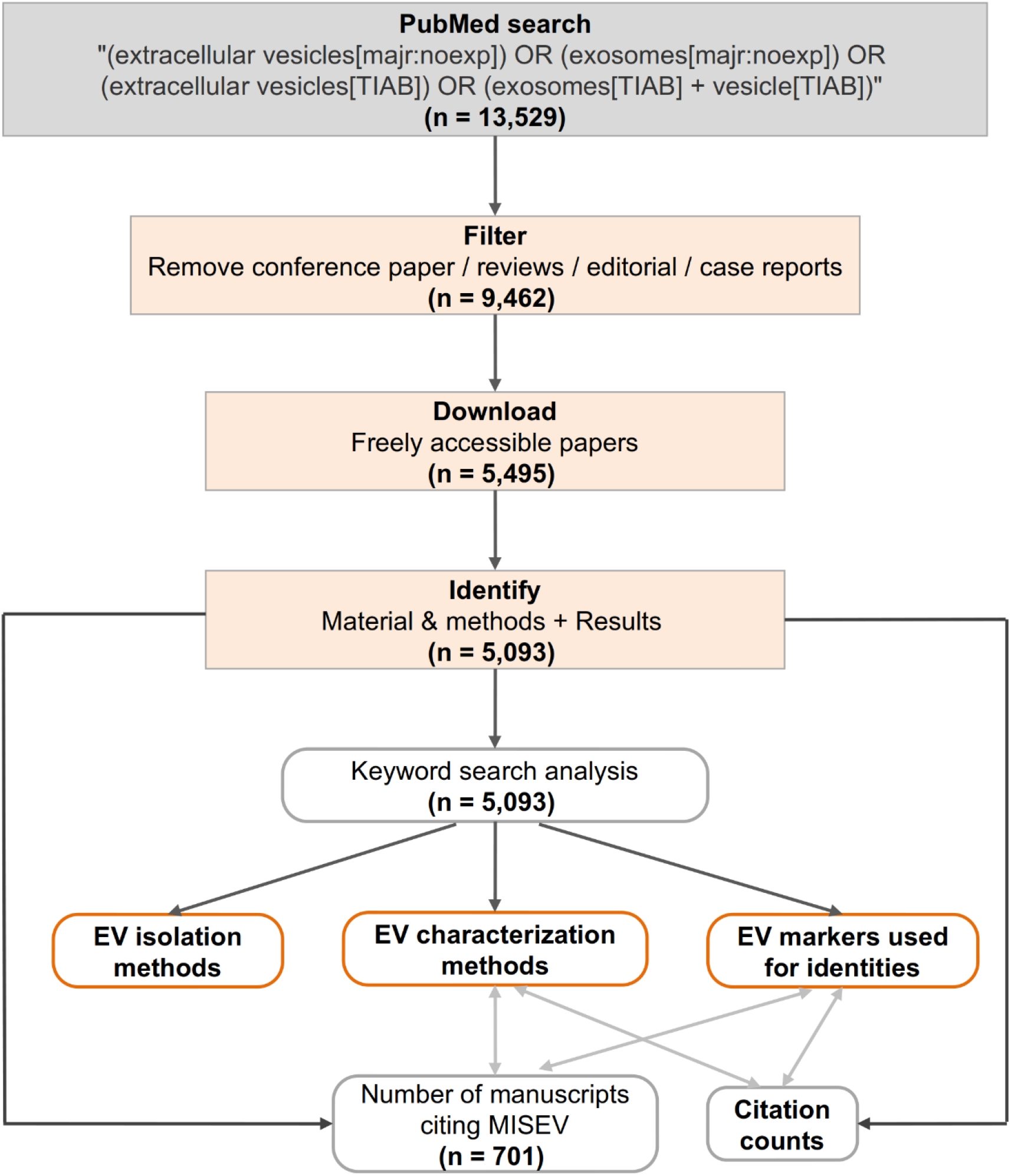
Dataset selection and approach used to investigate the compliance of the EV field to MISEV guidelines. Research was conducted in PubMed (grey box) and then filtering steps were applied to keep unique records, remove conference / reviews / editorials / case reports. The keyword search was conducted in open access papers where material and methods and/or result sections were found. In parallel, we also extracted citation counts and the list of papers citing MISEV guidelines [1,2] for the 5,093 selected publications.

## Notes

### Competing Interest Statement

The authors have declared no competing interest.

## References

[1] J. Lötvall, A.F. Hill, F. Hochberg, E.I. Buzás, D. Di Vizio, C. Gardiner, Y.S. Gho, I. V. Kurochkin, S. Mathivanan, P. Quesenberry, S. Sahoo, H. Tahara, M.H. Wauben, K.W. Witwer, C. Théry, Minimal experimental requirements for definition of extracellular vesicles and their functions: A position statement from the International Society for Extracellular Vesicles, J. Extracell. Vesicles. (2014). https://doi.org/10.3402/jev.v3.26913.

[2] C. Théry, K.W. Witwer, E. Aikawa, M.J. Alcaraz, J.D. Anderson, R. Andriantsitohaina, A. Antoniou, T. Arab, F. Archer, G.K. Atkin-Smith, D.C. Ayre, J.M. Bach, D. Bachurski, H. Baharvand, L. Balaj, S. Baldacchino, N.N. Bauer, A.A. Baxter, M. Bebawy, C. Beckham, A. Bedina Zavec, A. Benmoussa, A.C. Berardi, P. Bergese, E. Bielska, C. Blenkiron, S. Bobis-Wozowicz, E. Boilard, W. Boireau, A. Bongiovanni, F.E. Borràs, S. Bosch, C.M. Boulanger, X. Breakefield, A.M. Breglio, M. Brennan, D.R. Brigstock, A. Brisson, M.L.D. Broekman, J.F. Bromberg, P. Bryl-Górecka, S. Buch, A.H. Buck, D. Burger, S. Busatto, D. Buschmann, B. Bussolati, E.I. Buzás, J.B. Byrd, G. Camussi, D.R.F. Carter, S. Caruso, L.W. Chamley, Y.T. Chang, A.D. Chaudhuri, C. Chen, S. Chen, L. Cheng, A.R. Chin, A. Clayton, S.P. Clerici, A. Cocks, E. Cocucci, R.J. Coffey, A. Cordeiro-da-Silva, Y. Couch, F.A.W. Coumans, B. Coyle, R. Crescitelli, M.F. Criado, C. D’Souza-Schorey, S. Das, P. de Candia, E.F. De Santana, O. De Wever, H.A. del Portillo, T. Demaret, S. Deville, A. Devitt, B. Dhondt, D. Di Vizio, L.C. Dieterich, V. Dolo, A.P. Dominguez Rubio, M. Dominici, M.R. Dourado, T.A.P. Driedonks, F. V. Duarte, H.M. Duncan, R.M. Eichenberger, K. Ekström, S. EL Andaloussi, C. Elie-Caille, U. Erdbrügger, J.M. Falcón-Pérez, F. Fatima, J.E. Fish, M. Flores-Bellver, A. Försönits, A. Frelet-Barrand, F. Fricke, G. Fuhrmann, S. Gabrielsson, A. Gámez-Valero, C. Gardiner, K. Gärtner, R. Gaudin, Y.S. Gho, B. Giebel, C. Gilbert, M. Gimona, I. Giusti, D.C.I. Goberdhan, A. Görgens, S.M. Gorski, D.W. Greening, J.C. Gross, A. Gualerzi, G.N. Gupta, D. Gustafson, A. Handberg, R.A. Haraszti, P. Harrison, H. Hegyesi, A. Hendrix, A.F. Hill, F.H. Hochberg, K.F. Hoffmann, B. Holder, H. Holthofer, B. Hosseinkhani, G. Hu, Y. Huang, V. Huber, S. Hunt, A.G.E. Ibrahim, T. Ikezu, J.M. Inal, M. Isin, A. Ivanova, H.K. Jackson, S. Jacobsen, S.M. Jay, M. Jayachandran, G. Jenster, L. Jiang, S.M. Johnson, J.C. Jones, A. Jong, T. Jovanovic-Talisman, S. Jung, R. Kalluri, S. ichi Kano, S. Kaur, Y. Kawamura, E.T. Keller, D. Khamari, E. Khomyakova, A. Khvorova, P. Kierulf, K.P. Kim, T. Kislinger, M. Klingeborn, D.J. Klinke, M. Kornek, M.M. Kosanović, Á.F. Kovács, E.M. Krämer-Albers, S. Krasemann, M. Krause, I. V. Kurochkin, G.D. Kusuma, S. Kuypers, S. Laitinen, S.M. Langevin, L.R. Languino, J. Lannigan, C. Lässer, L.C. Laurent, G. Lavieu, E. Lázaro-Ibáñez, S. Le Lay, M.S. Lee, Y.X.F. Lee, D.S. Lemos, M. Lenassi, A. Leszczynska, I.T.S. Li, K. Liao, S.F. Libregts, E. Ligeti, R. Lim, S.K. Lim, A. Linē, K. Linnemannstöns, A. Llorente, C.A. Lombard, M.J. Lorenowicz, Á.M. Lörincz, J. Lötvall, J. Lovett, M.C. Lowry, X. Loyer, Q. Lu, B. Lukomska, T.R. Lunavat, S.L.N. Maas, H. Malhi, A. Marcilla, J. Mariani, J. Mariscal, E.S. Martens-Uzunova, L. Martin-Jaular, M.C. Martinez, V.R. Martins, M. Mathieu, S. Mathivanan, M. Maugeri, L.K. McGinnis, M.J. McVey, D.G. Meckes, K.L. Meehan, I. Mertens, V.R. Minciacchi, A. Möller, M. Møller Jørgensen, A. Morales-Kastresana, J. Morhayim, F. Mullier, M. Muraca, L. Musante, V. Mussack, D.C. Muth, K.H. Myburgh, T. Najrana, M. Nawaz, I. Nazarenko, P. Nejsum, C. Neri, T. Neri, R. Nieuwland, L. Nimrichter, J.P. Nolan, E.N.M. Nolte-’t Hoen, N. Noren Hooten, L. O’Driscoll, T. O’Grady, A. O’Loghlen, T. Ochiya, M. Olivier, A. Ortiz, L.A. Ortiz, X. Osteikoetxea, O. Ostegaard, M. Ostrowski, J. Park, D.M. Pegtel, H. Peinado, F. Perut, M.W. Pfaffl, D.G. Phinney, B.C.H. Pieters, R.C. Pink, D.S. Pisetsky, E. Pogge von Strandmann, I. Polakovicova, I.K.H. Poon, B.H. Powell, I. Prada, L. Pulliam, P. Quesenberry, A. Radeghieri, R.L. Raffai, S. Raimondo, J. Rak, M.I. Ramirez, G. Raposo, M.S. Rayyan, N. Regev-Rudzki, F.L. Ricklefs, P.D. Robbins, D.D. Roberts, S.C. Rodrigues, E. Rohde, S. Rome, K.M.A. Rouschop, A. Rughetti, A.E. Russell, P. Saá, S. Sahoo, E. Salas-Huenuleo, C. Sánchez, J.A. Saugstad, M.J. Saul, R.M. Schiffelers, R. Schneider, T.H. Schøyen, A. Scott, E. Shahaj, S. Sharma, O. Shatnyeva, F. Shekari, G.V. Shelke, A.K. Shetty, K. Shiba, P.R.M. Siljander, A.M. Silva, A. Skowronek, O.L. Snyder, R.P. Soares, B.W. Sódar, C. Soekmadji, J. Sotillo, P.D. Stahl, W. Stoorvogel, S.L. Stott, E.F. Strasser, S. Swift, H. Tahara, M. Tewari, K. Timms, S. Tiwari, R. Tixeira, M. Tkach, W.S. Toh, R. Tomasini, A.C. Torrecilhas, J.P. Tosar, V. Toxavidis, L. Urbanelli, P. Vader, B.W.M. van Balkom, S.G. van der Grein, J. Van Deun, M.J.C. van Herwijnen, K. Van Keuren-Jensen, G. van Niel, M.E. van Royen, A.J. van Wijnen, M.H. Vasconcelos, I.J. Vechetti, T.D. Veit, L.J. Vella, É. Velot, F.J. Verweij, B. Vestad, J.L. Viñas, T. Visnovitz, K. V. Vukman, J. Wahlgren, D.C. Watson, M.H.M. Wauben, A. Weaver, J.P. Webber, V. Weber, A.M. Wehman, D.J. Weiss, J.A. Welsh, S. Wendt, A.M. Wheelock, Z. Wiener, L. Witte, J. Wolfram, A. Xagorari, P. Xander, J. Xu, X. Yan, M. Yáñez-Mó, H. Yin, Y. Yuana, V. Zappulli, J. Zarubova, V. Žėkas, J. ye Zhang, Z. Zhao, L. Zheng, A.R. Zheutlin, A.M. Zickler, P. Zimmermann, A.M. Zivkovic, D. Zocco, E.K. Zuba-Surma, Minimal information for studies of extracellular vesicles 2018 (MISEV2018): a position statement of the International Society for Extracellular Vesicles and update of the MISEV2014 guidelines, J. Extracell. Vesicles. (2018). https://doi.org/10.1080/20013078.2018.1535750.

[3] M. Colombo, G. Raposo, C. Théry, Biogenesis, secretion, and intercellular interactions of exosomes and other extracellular vesicles, Annu. Rev. Cell Dev. Biol. (2014). https://doi.org/10.1146/annurev-cellbio-101512-122326.

[4] C. Gardiner, D. Di Vizio, S. Sahoo, C. Théry, K.W. Witwer, M. Wauben, A.F. Hill, Techniques used for the isolation and characterization of extracellular vesicles: Results of a worldwide survey, J. Extracell. Vesicles. (2016). https://doi.org/10.3402/jev.v5.32945.

[5] M. Palviainen, M. Saraswat, Z. Varga, D. Kitka, M. Neuvonen, M. Puhka, S. Joenväärä, R. Renkonen, R. Nieuwland, M. Takatalo, P.R.M. Siljander, Extracellular vesicles from human plasma and serum are carriers of extravesicular cargo—Implications for biomarker discovery, PLoS One. (2020). https://doi.org/10.1371/journal.pone.0236439.

[6] G. Van Niel, G. D’Angelo, G. Raposo, Shedding light on the cell biology of extracellular vesicles, Nat. Rev. Mol. Cell Biol. (2018). https://doi.org/10.1038/nrm.2017.125.

[7] L. Margolis, Y. Sadovsky, The biology of extracellular vesicles: The known unknowns, PLoS Biol. (2019). https://doi.org/10.1371/journal.pbio.3000363.

[8] S. El Andaloussi, I. Mäger, X.O. Breakefield, M.J.A. Wood, Extracellular vesicles: Biology and emerging therapeutic opportunities, Nat. Rev. Drug Discov. (2013). https://doi.org/10.1038/nrd3978.

[9] L. Doyle, M. Wang, Overview of Extracellular Vesicles, Their Origin, Composition, Purpose, and Methods for Exosome Isolation and Analysis, Cells. (2019). https://doi.org/10.3390/cells8070727.

[10] C. Théry, S. Amigorena, G. Raposo, A. Clayton, Isolation and Characterization of Exosomes from Cell Culture Supernatants and Biological Fluids, Curr. Protoc. Cell Biol. (2006). https://doi.org/10.1002/0471143030.cb0322s30.

[11] T.E. Whittaker, A. Nagelkerke, V. Nele, U. Kauscher, M.M. Stevens, Experimental artefacts can lead to misattribution of bioactivity from soluble mesenchymal stem cell paracrine factors to extracellular vesicles, J. Extracell. Vesicles. (2020). https://doi.org/10.1080/20013078.2020.1807674.

[12] J.B. Simonsen, What are we looking at? Extracellular vesicles, lipoproteins, or both?, Circ. Res. (2017). https://doi.org/10.1161/CIRCRESAHA.117.311767.

[13] V.V.T. Nguyen, K.W. Witwer, M.C. Verhaar, D. Strunk, B.W.M. Balkom, Functional assays to assess the therapeutic potential of extracellular vesicles, J. Extracell. Vesicles. (2020). https://doi.org/10.1002/jev2.12033.

[14] J.B. Simonsen, R. Münter, Pay Attention to Biological Nanoparticles when Studying the Protein Corona on Nanomedicines, Angew. Chemie - Int. Ed. (2020). https://doi.org/10.1002/anie.202004611.

[15] J. Van Deun, P. Mestdagh, P. Agostinis, Ö. Akay, S. Anand, J. Anckaert, Z.A. Martinez, T. Baetens, E. Beghein, L. Bertier, G. Berx, J. Boere, S. Boukouris, M. Bremer, D. Buschmann, J.B. Byrd, C. Casert, L. Cheng, A. Cmoch, D. Daveloose, E. De Smedt, S. Demirsoy, V. Depoorter, B. Dhondt, T.A.P. Driedonks, A. Dudek, A. Elsharawy, I. Floris, A.D. Foers, K. Gärtner, A.D. Garg, E. Geeurickx, J. Gettemans, F. Ghazavi, B. Giebel, T.G. Kormelink, G. Hancock, H. Helsmoortel, A.F. Hill, V. Hyenne, H. Kalra, D. Kim, J. Kowal, S. Kraemer, P. Leidinger, C. Leonelli, Y. Liang, L. Lippens, S. Liu, A. Lo Cicero, S. Martin, S. Mathivanan, P. Mathiyalagan, T. Matusek, G. Milani, M. Monguió-Tortajada, L.M. Mus, D.C. Muth, A. Németh, E.N.M. Nolte-’T Hoen, L. O’Driscoll, R. Palmulli, M.W. Pfaffl, B. Primdal-Bengtson, E. Romano, Q. Rousseau, S. Sahoo, N. Sampaio, M. Samuel, B. Scicluna, B. Soen, A. Steels, J. V. Swinnen, M. Takatalo, S. Thaminy, C. Théry, J. Tulkens, I. Van Audenhove, S. Van Der Grein, A. Van Goethem, M.J. Van Herwijnen, G. Van Niel, N. Van Roy, A.R. Van Vliet, N. Vandamme, S. Vanhauwaert, G. Vergauwen, F. Verweij, A. Wallaert, M. Wauben, K.W. Witwer, M.I. Zonneveld, O. De Wever, J. Vandesompele, A. Hendrix, EV-TRACK: Transparent reporting and centralizing knowledge in extracellular vesicle research, Nat. Methods. (2017). https://doi.org/10.1038/nmeth.4185.

[16] E.F. Pettersen, T.D. Goddard, C.C. Huang, G.S. Couch, D.M. Greenblatt, E.C. Meng, T.E. Ferrin, UCSF Chimera - A visualization system for exploratory research and analysis, J. Comput. Chem. (2004). https://doi.org/10.1002/jcc.20084.

[17] F. Royo, C. Théry, J.M. Falcón-Pérez, R. Nieuwland, K.W. Witwer, Methods for Separation and Characterization of Extracellular Vesicles: Results of a Worldwide Survey Performed by the ISEV Rigor and Standardization Subcommittee, Cells. (2020). https://doi.org/10.3390/cells9091955.

[18] S. Kovalchik, RISMED: Download Content from NCBI Databases, (2019).

[19] N. Jahn, roadoi: Find Free Versions of Scholarly Publications via Unpaywall, (2019). https://cran.r-project.org/package=roadoi.

[20] B. LeBeau, pdfsearch: Search Tools for PDF Files, J. Open Source Softw. (2018). https://doi.org/10.21105/joss.00668.

[21] S. Chamberlain, H. Zhu, N. Jahn, C. Boettiger, K. Ram, Client for Various “CrossRef” “APIs,” (2020).

[22] A. Tieu, M.M. Lalu, M. Slobodian, C. Gnyra, D.A. Fergusson, J. Montroy, D. Burger, D.J. Stewart, D.S. Allan, An Analysis of Mesenchymal Stem Cell-Derived Extracellular Vesicles for Preclinical Use, ACS Nano. (2020). https://doi.org/10.1021/acsnano.0c01363.

